# The Corrected Oxygenation Gradient: A Unified Index of Gas Exchange, Tissue Extraction, and Ventilatory Adequacy

**DOI:** 10.1101/2025.06.30.662232

**Authors:** Fardad Moshtagh Sisan, Christina Joya, Austin Situ, Scott Silver, Syed Akbarullah

## Abstract

Traditional gas exchange metrics, such as the alveolar-arterial (A–a) gradient (ΔP), fail to account for systemic oxygen extraction or ventilatory adequacy, thereby limiting their diagnostic precision in complex respiratory or circulatory pathologies. We propose and derive the corrected oxygenation gradient (COG), a novel formula that incorporates the A–a gradient, systemic oxygen extraction (CvO_2_/CaO_2_), and ventilatory adequacy (PaCO_2_ normalization). Complete mathematical proofs of continuity, differentiability, and boundary behaviors are provided. Simulations across physiological and pathological ranges illustrate logical behavior. COG appropriately scales with hypercapnia and tissue hypoxia, offering a single index sensitive to integrated oxygenation status. COG represents a theoretically rigorous and physiologically intuitive tool for characterizing gas exchange inefficiency.

## Introduction

Gas exchange assessment typically relies on the alveolar-arterial (A–a) oxygen gradient (ΔP), which is calculated as the difference between the estimated alveolar oxygen tension and the measured arterial oxygen tension [1]. However, the A–a gradient primarily reflects diffusion defects and ventilation-perfusion mismatch at the alveolar level. It fails to incorporate systemic oxygen extraction by tissues and changes in ventilatory effectiveness (e.g., hypercapnia) [2,3]. These limitations become critical in states such as septic shock, cardiopulmonary failure, or mixed respiratory-metabolic pathologies, where perfusion, extraction, and ventilation simultaneously affect gas exchange. We hypothesize that a combined index, the corrected oxygenation gradient (COG), can better quantify integrated pulmonary and systemic gas exchange.

## Materials and Methods

### Systemic Oxygen Extraction Correction

Systemic oxygen extraction reduces venous oxygen content (CvO_2_) relative to arterial oxygen content (CaO_2_). Systemic oxygen extraction is fundamentally defined as the ratio of CvO_2_ to arterial CaO_2_. For oxygen extraction correction, we define the extraction correction efficiency (ECE) as:

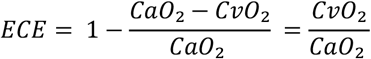

This ratio intuitively represents systemic tissue oxygen extraction: minimal extraction yields ECE=1 (CvO_2_≈CaO_2_), maximal extraction yields ECE=0 (CvO_2_≈0).

### Ventilatory Correction Factor (CCF) Derivation

For ventilatory correction based on measured arterial carbon dioxide tension (PaCO_2_), we defined a simple linear correction factor (CCF):

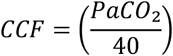

We chose a linear normalization around the baseline PaCO_2_ value of 40 mmHg, representing normal ventilation, to ensure direct proportionality and straightforward interpretation. Mathematically, this linear form provides a continuous, differentiable, and easily interpretable scaling around the physiological normal. If PaCO_2_ = 40 mmHg, then CCF would equal 1. If PaCO_2_ is greater than 40 mmHg (seen in hypercapnia), then CCF would be greater than 1, and if PaCO_2_ is less than 40 mmHg (seen in hypocapnia), then CCF would be less than 1.

### Corrected Oxygen Gradient (COG) Derivation

Both ECE and CCF are continuous, differentiable, dimensionless, and physiologically bounded. Integrating the two correction factors with the traditional alveolar-arterial gradient (ΔP), we define the corrected oxygenation gradient (COG) as:

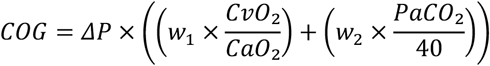

Weighting factors, are provisionally set equal to 1 each, pending empirical validation. The multiplicative integration reflects combined effects without altering dimensional integrity. Impaired alveolar-arterial oxygen transfer would increase ΔP, which then increases COG. Enhanced systemic extractions would decrease CvO_2_, reducing ECE and, thus, COG. Elevated PaCO_2_, such as in reduced ventilatory efficiency, would increase CCF and, therefore, COG.

### Computational Simulations

We conducted computational simulations to explore and visualize the behavior of COG across physiologically relevant ranges. The ΔP values ranged from 5 to 600 mmHg, representing conditions from normal pulmonary function to severe impairment in gas exchange. CvO_2_ ranged from 10 to 19 mL/dL, covering states of high systemic oxygen extraction to minimal extraction. PaCO_2_ values spanned from 20 to 80 mmHg, reflecting the full spectrum of ventilation status from marked hyperventilation to significant hypoventilation. Simulations and visualizations were performed using the Jupyter Notebook [4].

## Results

The COG demonstrated coherent and physiologically logical behavior across simulated conditions. A direct proportionality was observed between COG and PaCO_2_. As PaCO_2_ increased above the normal physiologic threshold of 40 mmHg, the COG rose accordingly, reflecting worsening ventilatory clearance. This linear amplification illustrated that hypercapnia exerts a measurable influence on effective oxygenation inefficiency, even when the primary diffusion defect remains constant (Figure 1).

**Figure 1.**
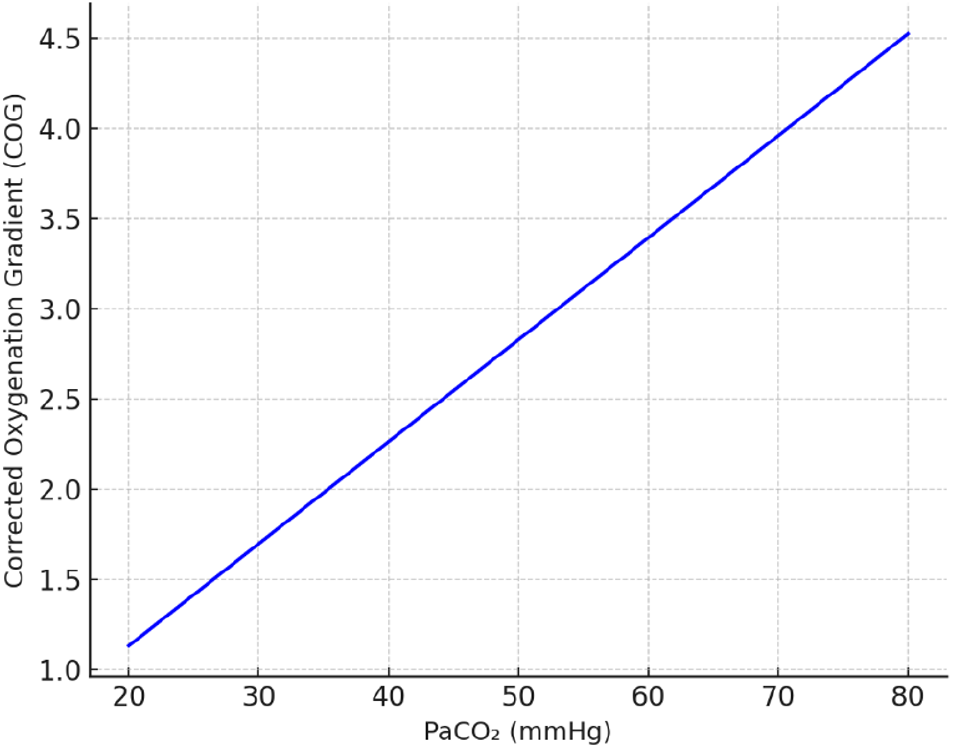
Relationship between corrected oxygenation gradient (COG) and arterial carbon dioxide tension (PaCO_2_). COG increases proportionally as PaCO_2_ rises above the physiologic baseline of 40 mmHg.

Similarly, as CvO_2_ decreased — indicating higher systemic tissue extraction — the COG declined proportionally (Figure 2). This behavior confirmed the model’s sensitivity to systemic oxygen utilization: cases with higher tissue oxygen extraction exhibited a protective effect by mitigating the apparent inefficiency of pulmonary gas exchange. Finally, a two-dimensional heatmap analysis integrating arterial oxygen saturation and PaCO_2_ demonstrated synergistic effects. Scenarios of combined low arterial oxygen saturation and elevated PaCO_2_ produced the highest COG values, indicating the most severe global gas exchange dysfunction (Figure 3). Conversely, states of preserved SpO_2_ and normocapnia maintained the lowest COG values, even with moderate diffusion gradients.

**Figure 2.**
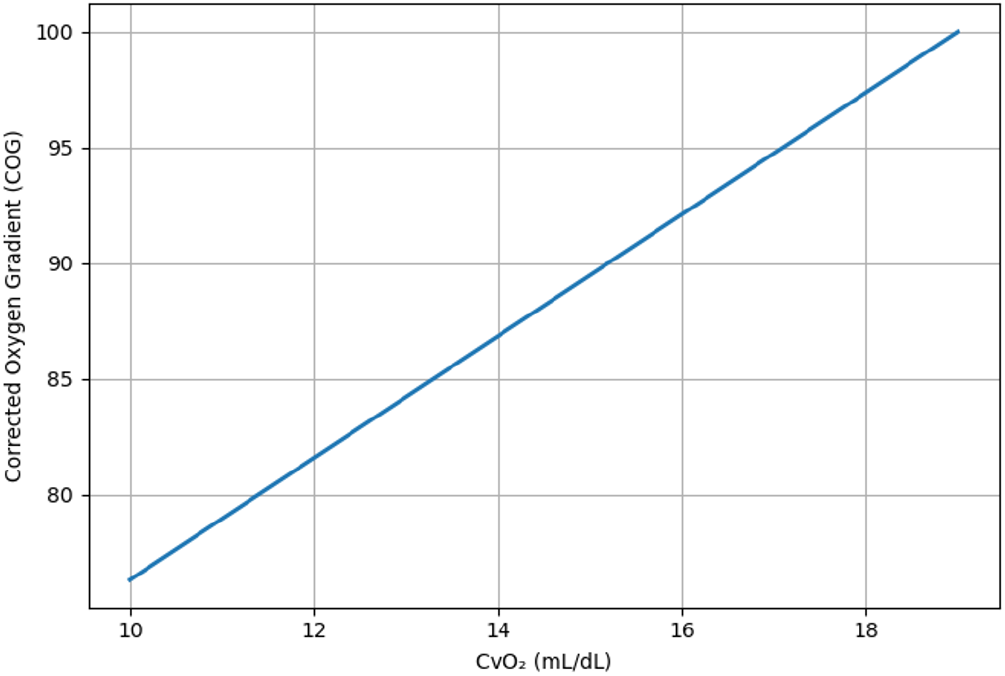
Relationship between corrected oxygenation gradient (COG) and venous oxygen content (CvO_2_). COG declines as CvO_2_ decreases, reflecting increased systemic tissue oxygen extraction.

**Figure 3.**
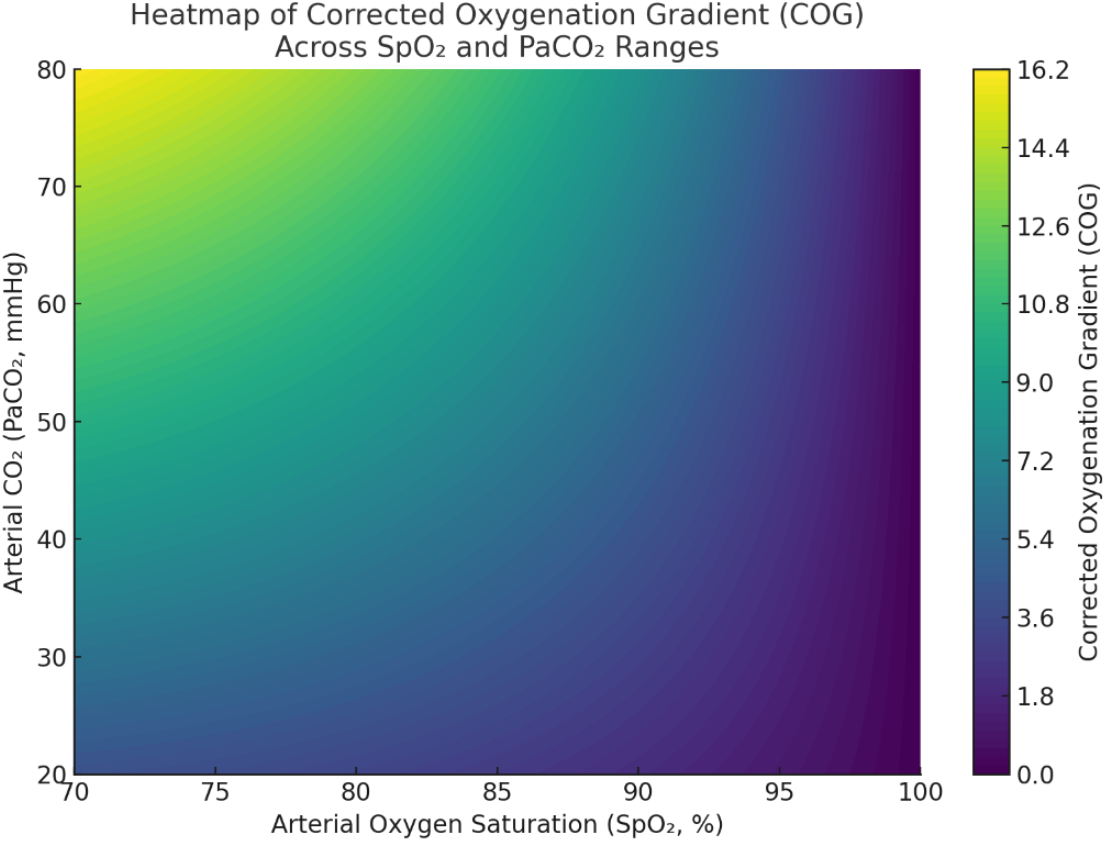
Heatmap illustrating the behavior of the corrected oxygenation gradient (COG) across a range of arterial oxygen saturations (SpO_2_) and arterial carbon dioxide levels (PaCO_2_). The COG increases sharply when SpO_2_ is low and PaCO_2_ is elevated.

## Discussion

The COG was developed to address limitations in traditional gas exchange assessments, particularly the A–a gradient. While the A–a gradient remains a valuable measure of pulmonary diffusion inefficiency, it fails to account for systemic oxygen extraction and ventilatory clearance, both of which profoundly influence oxygen delivery to tissues. Our results demonstrate that the COG integrates these critical factors into a single coherent index. The observed behavior of the COG aligns with physiological expectations. Rising PaCO_2_, representing impaired alveolar ventilation, proportionally amplified the COG, consistent with prior studies showing that hypercapnia worsens gas exchange efficiency independently of alveolar diffusion [1]. Similarly, decreases in CvO_2_—reflecting greater systemic tissue extraction—led to reduced COG values, reinforcing the concept that efficient tissue utilization of oxygen can partially compensate for upstream pulmonary inefficiency, as suggested in studies of systemic shock physiology [2,3]. Notably, the heatmap analysis revealed that concurrent hypoxemia and hypercapnia synergistically worsened the COG values. This finding supports clinical observations that scenarios with combined respiratory and metabolic failures exhibit more profound physiological compromise than those with isolated dysfunctions. By incorporating ventilatory status and tissue oxygen utilization into the evaluation of gas exchange, COG provides a more complete picture of patient physiology. It offers the potential to improve risk stratification, guide ventilatory management strategies, and serve as a prognostic marker in conditions ranging from septic shock to mixed respiratory failure. Future research directions include the clinical validation of the COG in prospective cohorts, correlation with clinical outcomes such as the need for mechanical ventilation, intensive care unit admission, and mortality, as well as exploration of its behavior in specific populations, including patients with chronic obstructive pulmonary disease, pulmonary hypertension, and heart failure.

This study has several limitations inherent to the development of theoretical models. First, the COG was derived and validated through mathematical analysis and simulation rather than direct clinical data. Although the model is physiologically grounded, prospective validation against real-world patient outcomes is necessary to confirm its predictive utility. Second, the calculations of CaO_2_ and CvO_2_ assume steady-state conditions and normal hemoglobin binding dynamics, which may not hold in conditions such as severe anemia, carbon monoxide poisoning, or methemoglobinemia. Third, ventilatory correction relies on PaCO_2_ as a surrogate for alveolar ventilation efficiency; however, factors such as metabolic acidosis, ventilator settings, or altered CO_2_ production could confound this relationship in complex clinical settings. Finally, while the correction factors were designed to be dimensionless and physiologically bounded, extreme pathologic states outside normal physiological ranges were not explicitly modeled. Further research should evaluate the impact of these assumptions and explore adaptations of COG for broader clinical scenarios and determining weighting factor for ECE and CCF.

## Conclusions

The COG provides a novel, mathematically validated and physiologically grounded, integrates pulmonary diffusion inefficiency, systemic oxygen extraction, and ventilatory adequacy into a comprehensive clinical tool. Preliminary simulations demonstrate robust physiological coherence across a broad spectrum of clinical scenarios. Prospective clinical validation and refinement of weighting factors will further optimize COG’s clinical utility, enhancing diagnostic precision and therapeutic guidance in critically ill patients.

## Notes

### Competing Interest Statement

The authors have declared no competing interest.

